# In vitro reconstitution of hippocampal cell assemblies

**DOI:** 10.64898/2025.12.19.695496

**Authors:** Cheng Gong, Aniekan Umoren, Alia Alameri, Estanislao Daniel De La Cruz, Bas Lendemeijer, Yannan Chen, Joseph A. Gogos, Steven A. Kushner, Kam W. Leong, Raju Tomer

## Abstract

Three-dimensional neuronal cultures, such as brain organoids, have proven effective for *in vitro* modeling of brain development and disease. However, it remains unclear whether these systems can intrinsically generate the spatiotemporal network motifs that support cognitive function, particularly given their lack of sensory input. To address this, we developed a 3D long-range modular neuronal network approach to test whether dissociated embryonic hippocampal cells can self-organize into neuronal manifolds capable of generating network dynamics analogous to the *in vivo* hippocampal cell assemblies implicated in cognitive function. Using all-optical interrogation, we demonstrate *in vitro* recapitulation of cell assemblies and their hallmark features, including recurring sequential activation, hierarchical chaining, multi-day stability, resilience to perturbation, and attractor-like pattern completion. We further investigated the effects of acute ketamine exposure on these assemblies, revealing network-wide reconfiguration reminiscent of *in vivo* responses. Finally, we show that human iPSC-derived neurons can similarly self-organize into modular network architectures. Together, these findings are consistent with an intrinsically “inside-out” view of the brain, rather than a *tabula rasa* model in which sensory input is required for the emergence of cognitive network motifs, and open new avenues for *in vitro* modeling of network-level mechanisms relevant to neuropsychiatric disorders.

## INTRODUCTION

Three-dimensional (3D) microphysiological systems (MPS), including brain organoids and assembloids, have emerged as effective *in vitro* platforms for modeling human neurodevelopment and diseases^1–4^. By partially recapitulating cytoarchitecture, cell-type diversity, and cell–cell interactions, iPSC-derived preparations have enabled mechanistic insights into disorders ranging from microcephaly^3^ to autism spectrum disorder^5^. However, despite their growing sophistication^2^, it remains unknown whether such systems can capture the spatiotemporal network dynamics, such as sequential activation, replay, metastable ensembles, and attractor-like computation, that underlie higher-order cognition. These motifs are central to neuropsychiatric disorders such as schizophrenia or depression, in which subtle disruptions in distributed network computation, rather than overt structural abnormalities, drive pathology^6–10^.

This limitation of current MPS intersects with a central question in systems neuroscience: do cognitive network motifs emerge primarily through sensory experience, or can they arise intrinsically from self-organizing neural manifolds? The “inside-out” framework^11^ and developmental studies of preconfigured hippocampal sequences^12,13^ suggest that the brain may possess endogenous attractor dynamics that precede and scaffold experience. Testing this hypothesis requires an *in vitro* system capable of spontaneously generating *bona fide* cell assembly-like patterns in the absence of any external stimuli.

We reasoned that dissociated cells from a defined brain structure might reconstitute its characteristic computational motifs if provided with an appropriate mesoscale architecture. We focused on the hippocampus, whose recurrent circuitry and topology are tightly linked to theoretical frameworks of memory encoding and spatial navigation^14–16^. *In vivo*, hippocampal cell assemblies - transient coalitions of neurons - exhibit recurring sequential activation, attractor-like pattern completion, multi-day stability with resilience to perturbation, and hierarchical chaining into “neural syntax”^17,18^. Demonstrating this repertoire *in vitro* would also shift MPS from models of cellular structure and excitability to models that implement the functional elements of cognition.

Previously, we introduced Modular Neuronal Networks (MoNNets^19^), in which dissociated embryonic hippocampal cells self-organize into interconnected spheroid modules on non-adhesive substrates. These networks exhibit hierarchical modular organization, diverse neuronal and glial cell types, and pharmacological responsiveness consistent with *in vivo* network physiology, and they have been used for quantitative modeling of schizophrenia-associated network dysfunctions. However, their limited spatial extent (∼10 units) constrained long-range activity flow and restricted the emergence of rich, multi-step assembly dynamics.

Here we develop Long-Range MoNNets (LR-MoNNets), macro-scale networks comprising hundreds of interconnected modules, and IncStim, a compact all-optical platform that enables high-speed, large field-of-view (>7 mm) Ca^2+^ imaging and patterned optogenetic stimulation entirely within a standard CO₂ incubator. Coupled with a graph traversal-based sequence motif detection pipeline, this system reveals spontaneously emerging Intrinsic Network Assemblies (INAs) that recapitulate key properties of hippocampal cell assemblies: recurrent sequential activation, hierarchical chaining, multi-day stability, resilience to perturbation, and attractor-like pattern completion upon partial optogenetic activation. We further show that low-dose ketamine, a rapid-acting antidepressant that has been shown to affect hippocampal ensembles and place-cell representations *in vivo*^20^, acutely reconfigures the INA repertoire. Finally, we demonstrate that human iPSC-derived neurons can self-organize into modular architectures that exhibit INA-like synchronized activity. Together, these findings suggest that elements of cognitive brain networks can emerge intrinsically from self-organizing, modular neural architectures independent of sensory experience, and position LR-MoNNets as a tractable *in vitro* system for dissecting network-level dysfunctions relevant to neuropsychiatric disease.

## RESULTS

### Development of Long-Range MoNNets and IncStim for all-optical interrogation

We previously introduced the MoNNet framework^19^, in which dissociated embryonic hippocampal cells spontaneously self-organize into interconnected modular spheroid units. Although these original MoNNets exhibited robust hierarchical organization, physiologically diverse cell populations, and gene expression signatures consistent with *in vivo* maturation, their limited spatial extent restricted the study of long-range activity propagation and complex, multi-step dynamics. We hypothesized that scaling these cultures to hundreds of interconnected modules, forming Long-Range MoNNets (LR-MoNNets), would permit the emergence and analysis of more elaborate sequential assembly dynamics (**Fig. 1a**, **Supplementary Video 1**). Increasing network size also facilitates macroscopic visualization of propagating activity patterns, mitigating the temporal constraints inherent to Ca²⁺ imaging.

**Figure 1.**
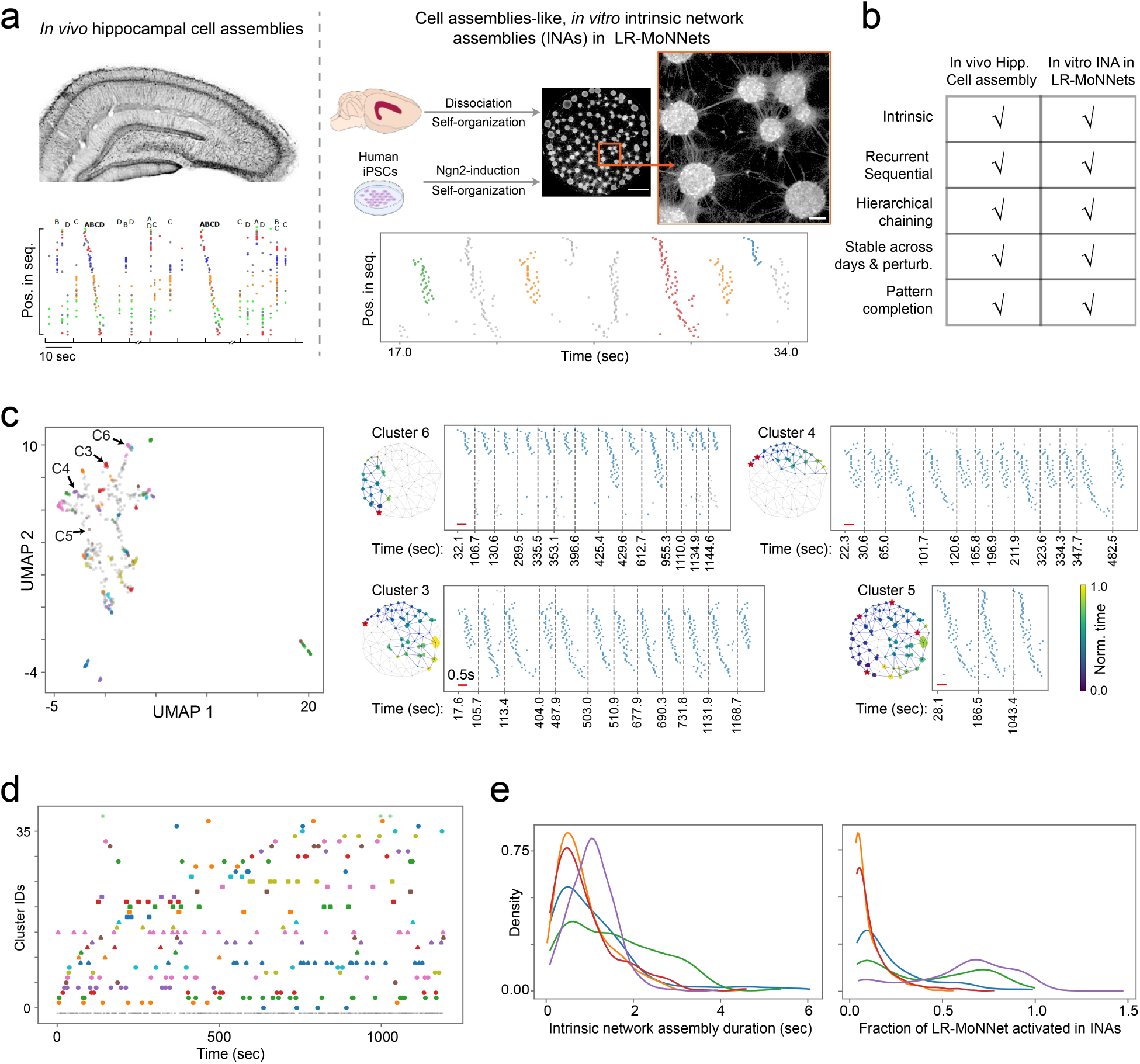
**In vitro Intrinsic Network Assemblies (INAs) in long-range MoNNets recapitulate hallmark features of hippocampal cell assemblies.** (a) *Left*: Schematic representation of *in vivo* hippocampal cell assemblies (adapted from Yuste *et al.*^17^). The plot below illustrates sequential neuronal activation within an assembly. *Right*: Schematic of the *in vitro* self-organization of dissociated hippocampal neurons (or human iPSC-derived neuronal progenitors) into Long-Range Modular Neuronal Networks (LR-MoNNets). The bottom plot shows representative sequential activity propagation within an Intrinsic Network Assembly (INA). See also **Supplementary Video 1** for Ca²⁺ imaging of two LR-MoNNet samples. Scale bars: 1 mm (full field); 100 μm (zoomed view). (b) Summary table comparing hallmark dynamic properties of *in vivo* hippocampal cell assemblies with those of LR-MoNNets reported in this study. (c) *Clustering and characterization of recurrent INA motifs*. *Left:* UMAP embedding of all detected INAs from a representative LR-MoNNet sample, with each point corresponding to an individual INA instance and colored by cluster identity. *Right:* Spatial activation maps and temporal dynamics of four representative INA clusters (C6, C4, C3, C5). For each cluster, the spatial motif is shown using a dark blue-to-yellow colormap indicating activation order, along with raster plots of all INA instances within that cluster. These examples demonstrate recurrent activation and hierarchical chaining (e.g., C6, C4, and C3 contributing to the composite sequence C5). All raster plots share a common y-axis ordering. See also **Supplementary Video 2** for an example of INA chaining. (d) Scatter plot illustrating spontaneous, recurring activation of INA clusters across time. Each dot represents a single INA event, colored by cluster membership. (e) *Left:* Distribution of INA durations, showing that assembly dynamics unfold on a timescale of seconds. *Right:* Distribution of the fraction of the LR-MoNNet recruited during individual INAs, demonstrating variability in assembly size across the repertoire. Data from five independent samples are shown in distinct colors.

To generate LR-MoNNets, we fabricated custom polydimethylsiloxane (PDMS) molds with 5.6-mm-diameter culture wells. Dissociated E18.5 mouse hippocampal cells were seeded at ∼550,000 cells per well, enabling the formation of large, three-dimensional modular architectures spanning a substantially greater area than in previous MoNNets. We implemented a two-stage viral transduction strategy to achieve all-optical compatibility: initial infection with pAAV.Syn.GCaMP6f (Addgene) for Ca²⁺ imaging, followed one week later by pAAV.Syn.ChrimsonR-tdTomato to enable red-shifted optogenetic activation. By the third week *in vitro*, these cultures displayed stable modular organization suitable for large-scale optical interrogation.

To record and manipulate activity within these macro-scale networks under stable physiological conditions, we engineered a compact, low-cost, large field-of-view (FOV) all-optical platform termed IncStim (**Supplementary Figure 1**, **Supplementary Table 1**). Built entirely from off-the-shelf components, IncStim fits inside a standard incubator, maintaining physiological temperature and CO₂ level to enable long-term imaging. The system provides a >7-mm FOV with cellular resolution (∼5 µm, **Supplementary Figure 1**) and supports arbitrary spatiotemporal optogenetic stimulation patterns. The imaging arm employs a 455-nm LED configured for uniform Köhler illumination for wide-field Ca²⁺ imaging. Note that the 455 nm imaging wavelength was specifically chosen to avoid any cross-excitation of ChrimsonR. The stimulation arm incorporates a compact RGB laser projector capable of projecting patterns from ∼15 µm to the full FOV. Importantly, we found that the red channel of the projector reliably drove ChrimsonR activation, enabling simultaneous imaging and targeted perturbation in an exceptionally compact form factor.

Together, the development of macro-scale LR-MoNNets and the engineering of the IncStim platform establish a fully integrated system for long-term, high-speed all-optical interrogation within physiologically controlled environments. This approach provides the experimental foundation necessary to systematically dissect large-scale network dynamics underlying complex computations.

### Detection and clustering of sequentially active Intrinsic Network Assemblies (INAs) in LR-MoNNets

Using the IncStim platform, we performed Ca²⁺ imaging of LR-MoNNets directly inside a cell-culture incubator to ensure stable physiological conditions. Images were acquired at 15 fps for all experiments. As shown in **Fig. 1c-d** and **Supplementary Videos 1-2**, LR-MoNNets exhibited richly structured sequential propagation of activation across the network. These spatiotemporal motifs, which we termed Intrinsic Network Assemblies (INAs), varied in configuration and engaged distinct combinations of interconnected modular units. We hypothesized that these INA motifs may represent *in vitro* analogs of hippocampal cell assemblies.

To test this hypothesis and to systematically analyze INA dynamics, we developed a custom computational pipeline for INA detection, clustering, and comparative analysis. Given the strongly synchronized intra-unit activity of each spheroid^19^ (**Supplementary Videos 1–2**) and the rapid propagation of activity across the macroscale network, Ca²⁺ signals from individual neurons within a spheroid appeared as a single, rapid burst. To overcome the temporal-resolution limitations inherent to Ca²⁺ indicators, we aggregated the activity of all neurons within each spheroid and treated each spheroid as a single activity node for subsequent motif analysis. This abstraction enabled robust delineation of inter-spheroid sequential dynamics.

The pipeline first computed the ΔF/F_o_ activity matrix by segmenting spheroids and extracting their mean fluorescence. Baseline fluorescence (F_o_) was estimated as the 20th percentile of camera background-corrected traces to provide a stable normalization. Spheroid bursts were then detected using an optimized threshold on the prominence of the ΔF/F_o_ trace peaks. To detect temporally isolated INAs, we developed a graph traversal-based algorithm. As a structural prior, the MoNNets were represented as graph using Delaunay triangulation, under the assumption that activity predominantly propagates through nearest-neighbor interactions - a premise supported by the observed spatiotemporal flow of activations. While long-range connections are present, this local-propagation prior enabled accurate segmentation of individual INA events. Building on this graph, a traversal-based algorithm was implemented to identify the complete INA repertoire within each recording.

To classify INA diversity and recurrence, we defined a custom distance metric capturing both the spatial and temporal components of INA structure (**Supplementary Figure 2**). This measure was used in a modified iterative k-means clustering procedure to identify cohesive sets of functionally similar INAs. For visualization, all INAs were embedded in UMAP (Uniform Manifold Approximation and Projection) space (**Fig. 1c**), revealing a diverse and well-separated landscape of spontaneously generated motifs. Importantly, clustering incorporated the full spatiotemporal trajectory of each INA sequence. Finally, to compare INA repertoires across experimental conditions, including pharmacological perturbations, we implemented an image-registration procedure that aligned spheroid positions across recordings, enabling direct unit-to-unit comparisons of assembly structure and dynamics.

### Intrinsic network assemblies recapitulate *in vivo* hippocampal cell assembly dynamics

Hippocampal cell assemblies are defined by hallmark population dynamics essential for spatial memory formation and cognitive processing. These canonical properties^17,18,21^ include: (1) recurrent activation and hierarchical chaining into structured sequences (“neural syntax”); (2) multi-day stability and resilience to network perturbation; and (3) attractor-like pattern completion that enables memory recall. We systematically evaluated whether the intrinsic network assemblies (INAs) identified in LR-MoNNets exhibit these defining *in vivo* features.

To assess the intrinsic recurrence and structural organization of INAs, we clustered all detected INA events and visualized their relationships using UMAP embedding. As shown in **Fig. 1c**, INAs displayed spontaneous and recurrent sequential activation, emerging in the absence of external inputs - consistent with “inside-out” models of intrinsic hippocampal circuit organization^11,13^. Moreover, we observed that smaller INA motifs frequently concatenated into larger, higher-order sequences (**Fig. 1c**, **Supplementary Video 2**), mirroring the hierarchical “neural syntax” proposed for *in vivo* cell assembly dynamics. Note that synchronized network activity within individual spheroids also represents smaller INAs, although they have been abstracted due to the technical limitations of Ca^2+^ indicators. Further, wave-duration analyses (**Fig. 1e**) revealed activity on timescales of seconds, comparable to the temporal structure of hippocampal assemblies in behaving animals^17^.

We next examined whether the INA repertoire maintains temporal stability, a hallmark of hippocampal cell assemblies that supports persistent memory representations. INAs detected in recordings separated by multiple days exhibited substantial cross-day correspondence (**Fig. 2a**), indicating preservation of assembly identity and suggesting a stable underlying network manifold. We further assessed INA robustness under perturbation, paralleling observations that hippocampal ensembles preserve integrity despite transient interference from other brain regions. Using an optogenetic perturbation paradigm in which ∼50% of the network was continuously stimulated for five minutes, we recorded activity before, during, and after the disturbance. As shown in **Fig. 2b** and **Supplementary Video 3**, a subset of INAs remained intact and re-emerged following stimulus cessation, demonstrating resilience and functional stabilization despite large-scale disruption.

**Figure 2.**
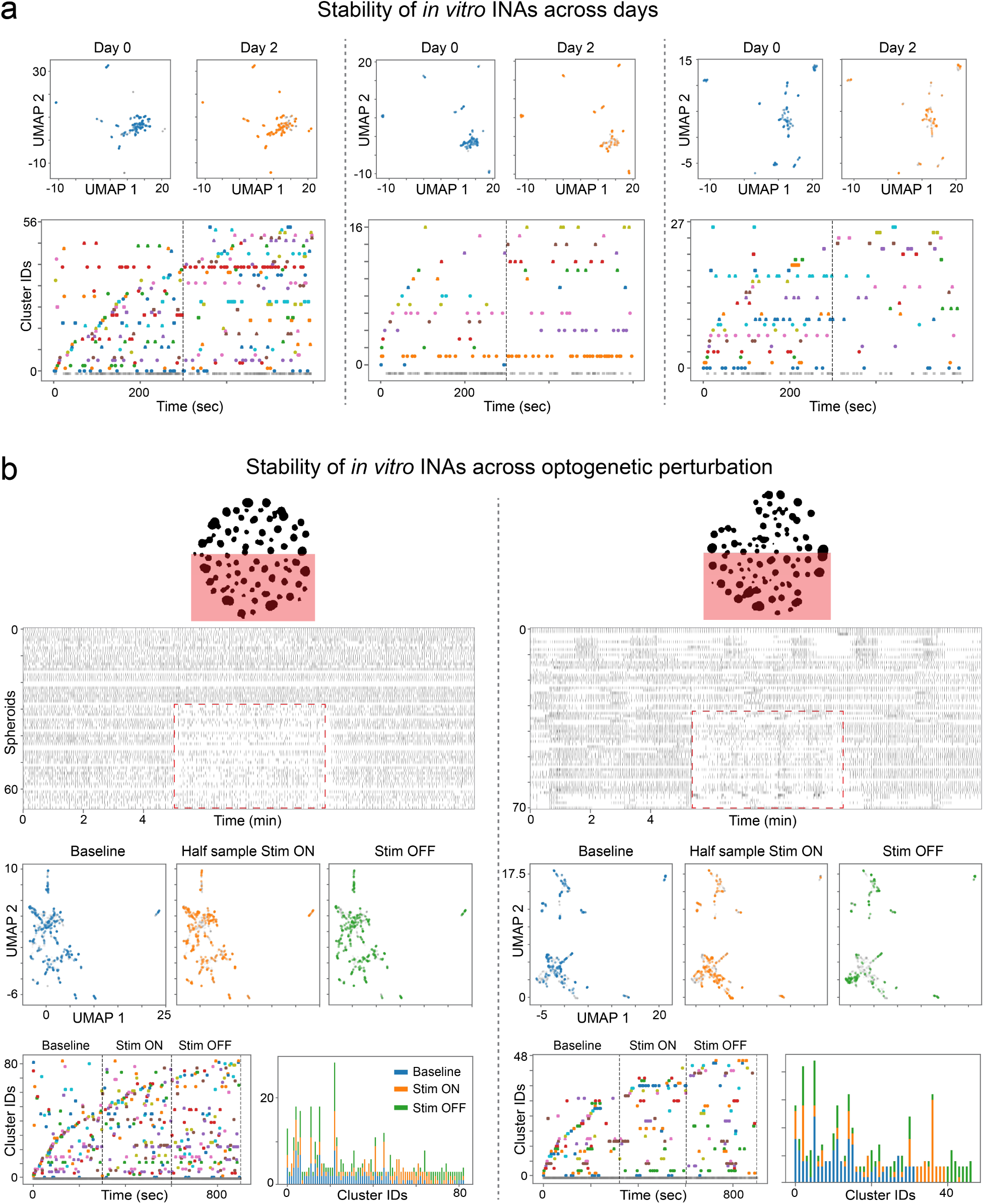
**Stability and resilience of *in vitro* Intrinsic Network Assemblies (INAs).** (a) *Multi-day stability of INA repertoires.* Three independent LR-MoNNet samples were imaged two days apart to assess temporal stability of INAs. *Top:* UMAP embeddings of INA instances detected on Day 0 and Day 2. The persistence of clusters in comparable regions of UMAP space indicates stable underlying functional network states across days. *Bottom:* Raster plots showing spontaneous recurrence of INAs (colored by cluster identity) on both days for each sample, demonstrating functional stabilization of specific assemblies over extended timescales. (b) *Resilience of INAs to optogenetic perturbation.* INA robustness was tested using a targeted, continuous optogenetic perturbation. *Top schematics:* Experimental paradigm in which approximately half of the LR-MoNNet was continuously stimulated (red shaded region). *Upper raster plots:* Network activity (spheroids vs. time, in minutes) recorded before (Baseline), during (Stim ON), and after (Stim OFF) perturbation for two representative samples. *Middle:* UMAP embeddings of INA instances detected during each phase. The reappearance of similar clusters in the recovery (Stim OFF) phase indicates preservation of the core INA repertoire despite large-scale disruption. *Lower:* Raster plots (cluster identity vs. time) and bar graphs quantifying INA recurrence frequency and cluster composition across phases. These analyses show that a subset of INAs survives the perturbation and reliably re-emerges during recovery, indicating functional resilience. See also **Supplementary Video 3** for Ca²⁺ imaging during all three phases.

Finally, we tested whether INAs implement pattern completion, a defining computation of hippocampal attractor networks in which partial cues reinstate full cell-assembly activity. We first identified robust spontaneous INAs and their corresponding temporal initiating units (**Fig. 3a**, top). Using the IncStim platform’s programmable patterned illumination, we then selectively activated ChrimsonR in these single initiating spheroids (indicated by red ROIs in **Fig. 3a**). As demonstrated by the graph-based activation maps and time-aligned raster plots (**Fig. 3b**), this partial input reliably reinstated the complete multi-unit INA trajectory. This reinstatement was robust across repeated trials and observed in independent samples (**Fig. 3**, **Supplementary Videos 4-5**). Together, these results indicate that LR-MoNNets perform an attractor-like computation, whereby the network state evolves from an incomplete cue toward the closest stored ensemble pattern, functionally analogous to hippocampal memory-retrieval dynamics.

**Figure 3.**
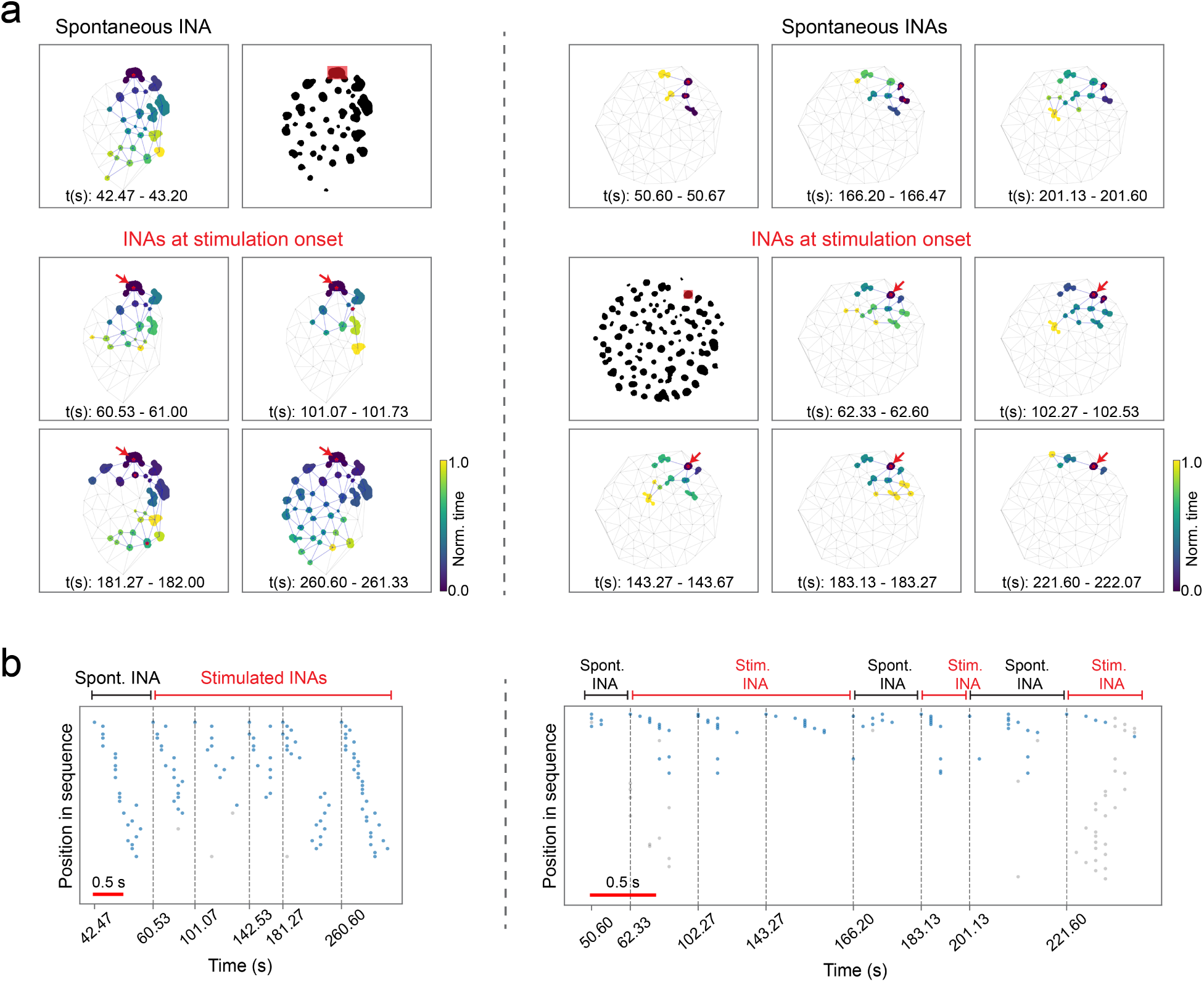
Pattern completion of INAs upon partial optogenetic activation. **(a)** Representative examples from two independent LR-MoNNet samples (separated by the vertical dashed line). *Top row*: Spontaneously occurring intrinsic network assemblies (INAs). *Bottom rows*: The corresponding INA reliably re-evoked by optogenetically stimulating a single modular unit. Network activity is visualized on the Delaunay graph of spheroid centroids, with nodes colored by normalized activation time (early-to-late). The optogenetic stimulation target is indicated by a red rectangle in the binary spheroid masks and by a red arrow in the sequence maps. The specific time window (t(s)) of each INA instance is listed at the bottom of each panel. **(b)** Raster representations of sequential activation for the examples in (a). Each dot represents the peak burst time of a spheroid, ordered by their position in the spontaneous sequence. The timelines illustrate that brief activation of a single initiating unit (red lines) consistently reinstates the full multi-unit INA trajectory across repeated trials. Timestamps on the x-axis correspond to the start times of the specific instances visualized in (a). Scale bar: 0.5 s. See also **Supplementary Videos 4-5** for these two representative examples.

Across all three defining dimensions - intrinsic recurrence and hierarchical syntax, multi-day stability and perturbation resilience, and attractor-like pattern completion - INAs in LR-MoNNets exhibit striking convergence with the established properties of hippocampal cell assemblies. These findings demonstrate that dissociated hippocampal networks, when reconstituted into long-range modular architectures, self-organize into functionally mature assemblies that recapitulate network elements of the hippocampal memory system.

### Acute ketamine exposure selectively reconfigures the INA repertoire in LR-MoNNets

To evaluate the translational utility of the LR-MoNNet platform for pharmacological investigation, we examined the acute effects of ketamine on intrinsic network assemblies (INAs). Ketamine, a dissociative anesthetic, is known to affect neuronal ensembles *in vivo*, and studies in rodents have demonstrated its inhibitory influence on place cells - key constituents of hippocampal cell assemblies^20^. Through concentration titration, we identified a low dose (250 nM) that induced measurable changes in network dynamics without causing global silencing and allowed rapid recovery upon media (-Ket) change (**Supplementary Video 6**), consistent with the transient nature of its clinical effects. Our experimental design included recordings from both control (H₂O) and ketamine-treated samples, each beginning with a 5-min baseline Ca²⁺ imaging period. Ketamine was then added to the medium, followed by immediate imaging (0 hr) and an additional recording one hour later (1 hr). All datasets were carefully registered to enable direct, unit-level comparisons of INA repertoires between baseline and treatment (or control) conditions.

As shown in **Fig. 4** and **Supplementary Video 6**, acute low-dose ketamine induced a pronounced reconfiguration of the INA repertoire. A substantial subset of previously active INAs became silenced or exhibited markedly reduced activation frequency, whereas others remained active. This demonstrates that ketamine does not merely suppress overall excitability but actively shifts the network into a distinct functional state, reorganizing the ensemble structure. Quantitative analysis across samples revealed several underlying physiological changes (**Fig. 4b**). Ketamine exposure significantly reduced mean activity rate and decreased both INA counts and INA durations. In parallel, inter-burst intervals increased, indicating slower, less frequent network activation. Notably, ΔF/F_o_ amplitude remained statistically unchanged. Critically, we observed a significant reduction in INA size (i.e., the number of participating units), suggesting that ketamine preferentially disrupts weaker functional linkages required for full propagation of sequential activity patterns.

**Figure 4.**
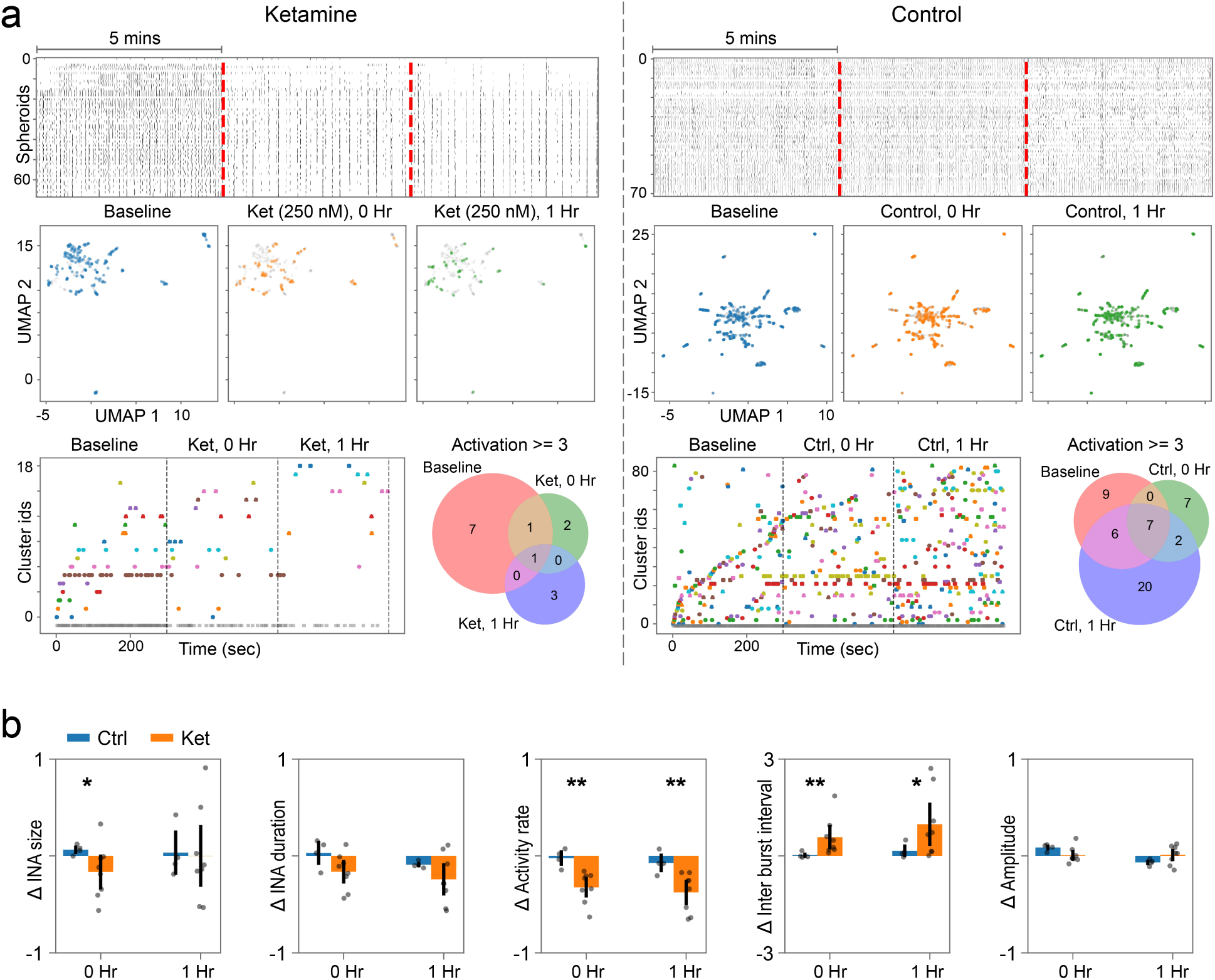
**Acute Ketamine exposure selectively reconfigures the INA repertoire in LR-MoNNets.** (a) *Dynamic reconfiguration of the INA repertoire following acute ketamine exposure. Top:* Raster plots showing spontaneous network activity in a representative LR-MoNNet treated with ketamine (250 nM) and a matched control (H₂O-treated) sample. Recordings were acquired at baseline, immediately after treatment (0 hr), and one hour later (1 hr). Ketamine treatment leads to a marked reduction in the frequency and density of network activation bursts relative to control. *Middle:* UMAP embeddings of detected INA instances for ketamine-treated and control samples across the three time points (Baseline, 0 hr, 1 hr). Ketamine exposure induces a pronounced shift and reduction in the number of distinct INAs, consistent with a reorganization of the functional network state. *Bottom:* Raster plots illustrating recurrent activation of INA clusters across conditions. Venn diagrams quantify the overlap of clusters exhibiting three or more activation events at each time point, highlighting the loss of pre-existing INAs following ketamine exposure. See also **Supplementary Video 6** for Ca²⁺ imaging at baseline, immediately after ketamine addition, one hour later, immediately after media replacement (−Ket), and one hour thereafter. (b) *Quantitative analysis of ketamine-induced changes in network dynamics.* Bar graphs show normalized changes in key INA metrics relative to baseline for ketamine-treated (Ket) and control (Ctrl) samples at 0 hr and 1 hr post-treatment. Acute exposure to 250 nM ketamine significantly reduced INA size and overall activity rate, while increasing inter-burst intervals, compared to control at both time points. INA duration showed a downward trend but did not reach statistical significance, and ΔF/F_o_ amplitude remained unchanged. Data are presented as mean ± SEM with individual sample values overlaid. Statistical significance is indicated by asterisks (*P < 0.05; **P < 0.01).

Together, these results demonstrate that acute ketamine exposure drives a dynamic reconfiguration of INA repertoires, characterized by reduced assembly size and diminished activation frequency. These findings align with *in vivo* observations of ketamine-induced inhibition and destabilization of hippocampal place cells and assemblies^20^, thereby validating the utility of LR-MoNNet/INA platform for probing of network-level mechanisms underlying psychopharmacological effects.

### Generation of human iPSC-derived, self-organized, modular neuronal networks

The successful recapitulation of complex, self-organizing, hippocampal cell assembly-like INAs in the mouse LR-MoNNet system motivated us to extend this framework to human cells. Because sequential cell assemblies are thought to represent fundamental computational motifs throughout the central nervous system^17,18^, we hypothesized that human iPSC-derived forebrain neurons might similarly self-organize into modular networks capable of exhibiting INA-like, stable sequential dynamics.

To test this, we implemented a rapid and efficient differentiation pipeline (summarized in **Fig. 5a**) based on doxycycline-inducible Neurogenin-2 (*Ngn2*) overexpression^22,23^. We utilized a previously developed, stable iPSC line^23^ harboring the *Ngn2* gene and introduced the calcium indicator GCaMP6f (driven by synapsin promoter) via lentiviral transduction at the pluripotent stage to ensure uniform expression across the resulting network. Following iPSC replating, *Ngn2* expression was induced via doxycycline to initiate rapid neurogenic differentiation.

**Figure 5.**
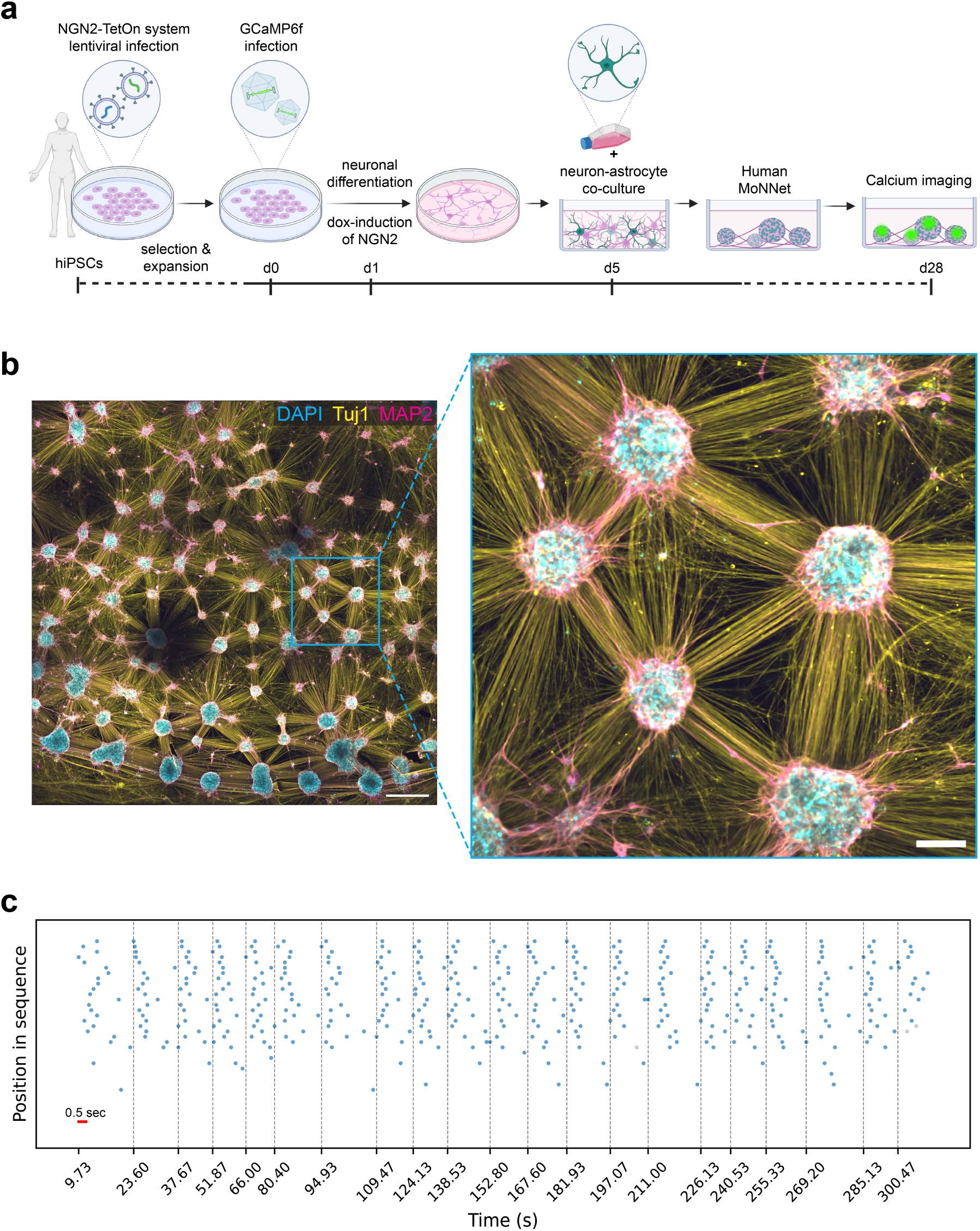
**Translation to human iPSC-derived, self-organized modular neuronal networks.** (a) *Generation of human long-range Modular Neuronal Networks (MoNNets).* Schematic of the differentiation and self-organization workflow. Human iPSCs (hiPSCs) harboring an NGN2-TetOn system were used for doxycycline-inducible neuronal differentiation. Cells were transduced with a lentiviral vector expressing the Ca²⁺ indicator GCaMP6f, followed by NGN2 induction. Differentiated neurons were subsequently co-cultured with astrocytes to promote maturation and spontaneous self-organization into modular human MoNNets, which were then used for Ca²⁺ imaging. (b) *Structural organization of human MoNNets.* Immunofluorescence images illustrating the self-organized modular architecture. *Left:* Wide-field view showing interconnected spheroid-like units distributed across the network. *Right:* Higher-magnification view highlighting local modular structure and interconnections. Cells are stained with DAPI (cyan), TUJ1 (yellow), and MAP2 (magenta), confirming neuronal identity and the formation of robust, interconnected modules. Scale bars: 500 µm (left), 100 µm (right). (c) *Spontaneous sequential Ca²⁺ activity in human MoNNets.* Representative ΔF/F_o_ traces recorded from human iPSC-derived LR-MoNNets, demonstrating intrinsically generated sequential activity patterns analogous to the INAs observed in mouse LR-MoNNets.

A critical determinant of modular self-organization was the introduction of glial support. On day 5 of differentiation, we introduced primary human astrocytes to the nascent neuronal cultures at a 1:1 ratio. This co-culture strategy proved essential; while neurons cultured in isolation formed flat monolayers, the addition of astrocytes promoted the spontaneous aggregation of neurons into three-dimensional, interconnected spheroid-like modules reminiscent of those observed in mouse LR-MoNNets. We further optimized the physical substrate to support these macroscopic structures, finding that while Matrigel supported short-term assembly, PLO-laminin coatings were required to maintain the structural integrity of these networks for long-term recordings (>8 weeks). Immunocytochemistry performed on these mature co-cultures confirmed the generation of a defined neuro-glial architecture.

As shown in **Fig. 5b**, the human iPSC-derived cultures displayed clear modular organization with well-defined structural units, similar to the mouse LR-MoNNets. Ca²⁺ imaging of spontaneous activity (**Fig. 5c**) revealed that these modules did not fire stochastically but instead exhibited robust, synchronized, sequential activation patterns. These spatiotemporal dynamics resembled the INA motifs identified in the mouse LR-MoNNet preparations, indicating an intrinsic capacity for self-organization into modular, sequentially active networks, when provided with appropriate mesoscale architecture and glial support.

## DISCUSSION

A longstanding challenge in neuroscience has been the inability of *in vitro* systems to reproduce the complex spatiotemporal codes that underlie higher-order cognition, such as sequential activation, assembly formation, and replay. Here we demonstrate that dissociated embryonic hippocampal cells can self-organize into long-range modular neuronal networks that generate robust, cell assembly-like intrinsic network assemblies (INA). Using our compact IncStim platform together with a custom dynamic motif-detection computational pipeline, we show that INAs recapitulate the hallmark dynamics of *in vivo* hippocampal assemblies: recurrent activation, hierarchical chaining, multi-day stability, resilience to perturbation, and attractor-like pattern completion evoked by partial optogenetic cues (summarized in **Fig. 1a–b**). These results establish that the core computational primitives of hippocampal circuits are not exclusive to the intact brain but can emerge *in vitro* when the architecture provides appropriate scale, heterogeneity, and possibilities for self-organization.

Our findings also advance the longstanding debate surrounding the developmental origins of cognitive circuits. A central question in systems neuroscience is whether dynamic motifs such as “neural syntax” arise through sensory-driven learning or reflect intrinsic structure embedded within a pre-existing neural manifold. The emergence of robust sequential dynamics in our sensory-deprived system challenges the strictly constructivist view. Instead, our results support a model in which the hippocampus possesses a genetically specified scaffold for sequential organization that operates independently of external input. This aligns with the “pre-configured” hypothesis^11–13,21^, suggesting that the brain is not a *tabula rasa* but develops a dynamical manifold that is subsequently refined, rather than created, by experience to support information processing and memory indexing.

These observations carry evolutionary implications. The hippocampus is an ancient archicortical structure that relies on recurrent collaterals to implement auto-associative memory computations. The ability of dissociated neurons to reconstitute assembly-like dynamics suggests that the organizing rules of archicortical circuits are encoded at the cellular and meso-scale levels, independent of intact anatomical wiring. This raises the possibility that self-organized computation is a conserved property of ancient cortical architectures, positioning LR-MoNNets as a potentially generalizable template for modeling other recurrent neural systems.

From a translational perspective, our platform shifts pharmacological screening from coarse metrics such as cytotoxicity or firing rate toward dynamics-based biomarkers. We found that low-dose ketamine does not simply suppress activity but selectively reconfigures the INA repertoire, destabilizing some assemblies while sparing others. This is consistent with the “disconnection hypothesis” of psychosis and with ketamine’s dissociative mechanism, in which the coherence of cognitive ensembles is disrupted. INA stability therefore emerges as a nuanced functional readout for assessing compounds that modulate cognitive network motifs. Moreover, our demonstration that human iPSC-derived neurons self-organize into modular architectures and exhibit sequential activity patterns provides a foundation for future work aimed at patient-specific modeling of network-level dysfunctions.

There are several limitations to consider. The all-optical approach of the LR-MoNNet platform was designed to capture long-range, complex network dynamics at the systems level, but this comes at the cost of reduced temporal and cellular resolution. The inherently slow kinetics of Ca²⁺ indicators, combined with large field-of-view imaging, limit the resolution of fast activity propagation within individual spheroid modules at single-spike timescales; consequently, we likely underestimate both the diversity of INA motifs and the fine structure of intra-spheroid dynamics. Complementary approaches, such as microelectrode arrays, could improve temporal precision, though typically at the expense of full-system coverage. Additionally, while nearest-neighbor Delaunay triangulation provides robust INA event segmentation, this structural prior may fail to capture some of the long-range “shortcut” connections that contribute to propagation. Finally, the Ngn2-based induction strategy for human iPSC-derived neurons, even with astrocyte co-culture, may not recapitulate the full diversity of native neuronal and glial subtypes; future iterations will benefit from incorporating additional cell types, particularly inhibitory interneuron populations.

Overall, by shifting the focus from structural descriptions of static phenotypes to functional modeling of dynamic cognitive states, our approach opens new avenues for dissecting the circuit-level mechanisms underlying complex neuropsychiatric disorders and for evaluating therapeutic interventions within a physiologically relevant, computationally active model system.

## METHODS

### Experimental procedures

#### Fabrication of PDMS culture molds

Custom polydimethylsiloxane (PDMS) culture molds were fabricated to support Long-Range MoNNet formation. First, negative casts were 3D-printed using acrylonitrile butadiene styrene (ABS) on an Ultimaker 2+ printer. Fresh PDMS was prepared by mixing the silicone elastomer base and curing agent (Sylgard 184) at a ratio of 10:1 (w/w), followed by degassing in a vacuum chamber to remove air bubbles. The pre-polymer mixture was poured onto the custom casts, covered with a coverslip to ensure flatness, and cured at 75°C overnight. The resulting PDMS molds, containing wells with a diameter of 5.6 mm, were released from the casts and sterilized by immersion in 100% ethanol overnight prior to use.

#### Preparation of mouse LR-MoNNets

All experimentations and animal handling followed US National Institutes of Health guidelines. All procedures were approved by the Institutional Animal Care and Use Committees (IACUC, AC-AABL2553) of Columbia University. LR-MoNNets were generated from embryonic day 18.5 (E18.5) mouse hippocampal tissue. Pregnant female mice (Wild-type, CD1) were anesthetized with isoflurane and euthanized by cervical dislocation. Embryos were harvested and placed in cold Hanks’ Balanced Salt Solution (HBSS; Gibco). Hippocampi were dissected in ice-cold Hibernate-E medium (Gibco) and incubated in 0.25% Trypsin-EDTA (Gibco) at 37°C for 30 minutes. Digestion was terminated by the addition of quenching medium (Hibernate-E supplemented with FBS) containing DNAse I (1 μg/mL; Sigma) at room temperature. Mechanical dissociation was performed by repeated pipetting with fire-polished glass Pasteur pipettes until a single-cell suspension was obtained. Cell viability was verified using the Trypan Blue exclusion method. The suspension was centrifuged at 900 rpm for 10 minutes, and the supernatant was aspirated. The resulting pellet was resuspended in culture medium consisting of Neurobasal medium supplemented with 1X B27, 0.5 mM Glutamax, and 1% Penicillin/Streptomycin (all from Gibco). The cell density was quantified using a hemocytometer. Prior to seeding, the cell suspension was transduced with an adeno-associated virus (AAV) expressing the calcium indicator GCaMP6f under the control of the synapsin promoter (pAAV.Syn.GCaMP6f.WPRE.SV40; Addgene). Cells were seeded into the sterilized PDMS molds at a density of 550k per well. The suspension was allowed to settle for 45 minutes in a humidified incubator at 37°C to promote initial aggregation before the addition of 2 mL of culture medium. Cultures were maintained at 37°C in a humidified atmosphere containing 5% CO_2_. One week post-seeding, during a scheduled media change, cells were super-infected with pAAV-Syn-ChrimsonR-tdT (Addgene) to enable optogenetic manipulation via red shifted ChrimsonR. Viral expression was robustly visible two weeks post-infection. By the third week *in vitro*, cultures exhibited stable GCaMP6f and ChrimsonR expression and were used for all-optical interrogation experiments.

#### Preparation of Human LR-MoNNets and immunostaining

As summarized in **Fig. 5a**, differentiation of human induced pluripotent stem cells (hiPSCs) into excitatory neurons was achieved through overexpression of the neurogenic transcription factor Neurogenin-2 (*Ngn2*). Human hiPSCs harboring a doxycycline-inducible Ngn2 gene (dox-hiPSCs) were generated according to established methods and have been described previously^23^. Dox-hiPSCs were plated on Matrigel-coated dishes (Corning, 356230), expanded in StemFlex™ medium (Thermo Fisher Scientific, A3349401), and maintained at 37 °C with 5% CO₂. Upon reaching 60–70% confluence, cells were dissociated using Accutase® (Sigma, A6964), centrifuged, and prepared for replating onto poly-L-ornithine (PLO)-laminin- or Matrigel-coated dishes.

To enable downstream live Ca²⁺ imaging, dissociated cells were resuspended in viral infection medium consisting of StemFlex™ supplemented with 1% RevitaCell™ (Thermo Fisher Scientific, A2644501) and pAAV.Syn.GCaMP6f.WPRE.SV40 (Addgene, plasmid #100837). Replating onto PLO-laminin-coated dishes was used for long-term cultures (>8 weeks) due to higher coating stability, whereas Matrigel-coated dishes were used for short-term cultures (up to 5 weeks). Both coating conditions yielded self-organizing spherical structures with synchronous activity.

The following day, viral infection medium was replaced with StemFlex™ containing doxycycline (4 µg/mL) to initiate *Ngn2* induction. One day later, the medium was switched to neuronal induction medium composed of Advanced DMEM/F12 (Thermo Fisher Scientific, 12634010), 1% N-2 supplement (Thermo Fisher Scientific, 17502048), and freshly added growth factors supporting neuronal maintenance: brain-derived neurotrophic factor (BDNF; 10 ng/mL, PeproTech), neurotrophin-3 (NT-3; 10 ng/mL, PeproTech), laminin (1 µg/mL, Sigma, L2020), and doxycycline (4 µg/mL). Doxycycline, BDNF, NT-3, and laminin were maintained in the culture medium for the remainder of the differentiation period.

On day 3, the base medium was transitioned to neuronal maturation medium consisting of a 1:1 mixture of Neurobasal (Thermo Fisher Scientific, 21103049) and DMEM/F12 + GlutaMAX™ (Thermo Fisher Scientific, 10565018), supplemented with 2% B-27 supplement (Thermo Fisher Scientific, 12587010). All base media were supplemented with gentamicin (50 µg/mL, Thermo Fisher Scientific, 15710064) to prevent contamination.

Primary human astrocytes (HAs; ScienCell, 1800) were cultured separately on poly-L-lysine-coated dishes and maintained in Astrocyte Medium (ScienCell, 1801) according to the manufacturer’s instructions. On day 5, neuronal maturation medium was supplemented with cytosine β-D-arabinofuranoside (Ara-C; 2 µM), and HAs were trypsinized and added to the neuronal cultures at a 1:1 ratio. On day 7, Ara-C was withdrawn by refreshing the neuronal maturation medium. Thereafter, 50% medium changes were performed every two days to maintain co-cultures while minimizing mechanical disturbance.

For immunocytochemistry, human neuro-glial co-cultures were fixed in 4% formaldehyde in PBS for 15 min at room temperature and washed three times with PBS. Samples were blocked for 1 h at room temperature in blocking buffer containing 5% normal donkey serum (Jackson ImmunoResearch, 017-000-121), 0.05% bovine serum albumin (Fisher Scientific, BP9703-100), and 0.2% Triton™ X-100 (Sigma, X100). Primary antibodies, chicken anti-MAP2 (1:1500, Thermo Fisher Scientific, PA1-10005) and rabbit anti-TUJ1 (1:300, Cell Signaling Technology, 5568S), were diluted in blocking buffer and incubated for 1 h at room temperature. Samples were washed three times with PBS (5 min per wash) and incubated for 1 h at room temperature with secondary antibodies diluted in blocking buffer: donkey anti-chicken Alexa Fluor™ 488 (1 µg/mL, Jackson ImmunoResearch, 703-546-155), donkey anti-rabbit Alexa Fluor™ 594 (8 µg/mL, Thermo Fisher Scientific, A-21207), and Hoechst 33342 (1 µg/mL, Tocris Bioscience, 5117). Samples were then washed three times with PBS (5 min per wash).

### IncStim Development

The IncStim system was designed to integrate wide-field Ca^2+^ imaging and patterned optogenetic stimulation into a single, compact platform designed for operation within a standard CO_2_ incubator. The system provided a field of view greater than 7 mm and achieved a lateral resolution of 5 µm at the center of the field (**Supplementary Figure 1**), exceeding that of standard fluorescence microscopes and enabling simultaneous visualization and stimulation of large MoNNet samples. This configuration enables stable, long-term imaging of LR-MoNNet samples or other live cultures under physiologically relevant environmental conditions. The mechanical and optical architecture was constructed primarily using a Thorlabs cage system, resulting in an overall footprint of 6x6x18 cu. inch, allowing the device to fit entirely within the incubator space.

#### Optical Architecture

As described in **Supplementary Figure 1** (see **Supplementary Table 1** for detailed parts list), the imaging arm incorporated a blue-shifted LED light source (center wavelength 455 nm, bandwidth 18 nm, Thorlabs) to excite the GCaMP6f calcium indicator. Note that 455 nm was specifically chosen to avoid inadvertent activation of ChrimsonR. The illumination path consisted of an aspherical collimation lens, an adjustable diaphragm, a focusing lens, and a large-field-of-view objective. This setup was precisely configured to achieve Köhler illumination with an optimized output power of 0.4 mW at the sample plane. This arrangement ensured uniform, stable, and wide-area illumination essential for high-quality live imaging of large samples. The optogenetic stimulation arm utilized a portable three-color RGB laser projector (Ultimems) capable of delivering arbitrary spatially patterned light. The optical path employed a pair of scan lenses, a focusing lens, and a fold mirror, followed by a long-pass dichroic beam splitter (DBS) to accurately direct the patterned excitation light onto the sample. Samples were mounted on a long range manual XYZ translation stage (OWIS), facilitating fine positional alignment.

#### Optogenetic stimulation pattern generation

Patterned stimulation was controlled using a custom Python-based interface that supported user-defined illumination designs. The size of each stimulation element could be adjusted by specifying its height, width and coordinates, enabling single stimulation blocks ranging from 15 µm up to the full field-of-view. Multiple stimulation blocks could be defined independently, with each block’s position and dimensions set programmatically. Stimulation was delivered at 2 Hz with an on/off duty cycle of 10–90%, and the laser output was set to 0.05mW to achieve effective activation while minimizing photobleaching in MoNNet samples.

### Ca^2+^ imaging, optogenetic manipulations and ketamine treatment

#### General recording procedures

All LR-MoNNet samples were transferred into a Petri dish and placed inside the IncStim imaging incubator for a minimum of 2 hours to allow for temperature and gas (CO_2_) equilibration. After stabilization, the sample dish lid was removed, and the dish was gently positioned onto the sample holder. The XYZ translation stage was manually adjusted to bring the MoNNet structures into focus. All Ca^2+^ imaging were acquired at a frame rate of 15 frames per second (fps). For long-term recordings (spanning multiple days), the sample dish was removed after each imaging session and stored in the incubator with the lid closed until the next scheduled imaging cycle.

#### Optogenetic stimulation

Optogenetic stimulation was applied simultaneously with image acquisition and could be adjusted flexibly during recording. Before each recording, the green channel of the RGB laser projector was used to visually locate and define the Region of Interest (ROI) on the live image. The size and position of the stimulation pattern were defined using the custom software interface. Subsequently, the system was switched to the red channel (635 nm), which activates the red-shifted ChrimsonR. Crucially, the red excitation light was not detectable by the imaging camera due to the exclusion by the GCaMP emission band-pass filter. Recordings were then initiated, and patterned stimulation was applied concurrently according to the experimental requirements.

#### Ketamine treatment experiments

Paired recordings were performed for each pharmacological investigation, with one MoNNet sample serving as the vehicle control and the other treated with ketamine. Each experiment began with a 5-minute baseline recording. For the ketamine-treated samples, 4 µL of a 250 µM ketamine stock solution was gently dispensed onto the edge of the sample. Given that each dish contained 4 mL of culture medium, this resulted in a final working concentration of approximately 250 nM ketamine after mixing. For the control sample, an equal volume of H_2_O (Vehicle) was added. Follow-up recordings were acquired immediately (0 Hr) and 1 hour (1 Hr) after treatment. For the ketamine washout experiments, the medium in the treated samples was completely replenished with fresh culture medium. Activity was recorded immediately and 1 hour after the washout. All control samples were recorded at the same corresponding time points for comparison.

### Data analysis

#### Detection of Spheroid Bursts

We first computed the ΔF/F₀ activity matrix by segmenting spheroids and extracting their mean fluorescence signals. Baseline fluorescence (F₀) was estimated as the 20th percentile of the camera background-corrected fluorescence traces. Spheroid bursts were then detected using an optimized threshold on peak prominence, which quantifies how much a peak stands out relative to surrounding peaks in the ΔF/F₀ trace. When the signal-to-noise ratio was sufficient, peaks were classified as noise or signal based on their prominence. The procedure was as follows: (1) identify local maxima in each ΔF/F₀ trace and compute their prominences using the SciPy signal-processing module (v1.12.0) in Python; (2) generate an empirical distribution of peak prominences using Gaussian kernel density estimation, with the bandwidth manually optimized to ensure that the noise component was represented by a single mode; (3) compute the first and second derivatives of the empirical distribution and set the prominence threshold at the point where the first derivative approaches zero and the second derivative is non-negative; (4) define peak duration as the full width at 40% of the peak’s maximum height; (5) repeat steps 1-4 for each activity trace in the ΔF/F₀ matrix. We found that this procedure outperformed a conventional 2-standard deviation thresholding approach in detecting spheroid activity bursts.

#### Detection of INAs

To detect sequential activation events (INAs), we used Delaunay triangulation to construct a structural prior graph, motivated by the observation that activity in LR-MoNNets predominantly propagates via nearest-neighbor interactions. Although long-range connections are present in LR-MoNNets, this local-propagation assumption enabled accurate detection of individual INA events using a graph traversal-based breadth-first search (BFS) algorithm. Traversal was constrained to neighboring nodes in the Delaunay graph that were active within the onset-to-peak interval of the initiating spheroid. This approach allows detection of overlapping INAs, which were merged in a final post-processing step.

#### Clustering of INAs

To classify and quantify the diversity and recurrence of INAs, we defined a custom distance metric, based on Jaccard distance, that captures both the spatial and temporal components of INA structure (**Supplementary Figure 2**). This metric was used in a modified iterative *k*-means-like clustering procedure to identify cohesive sets of functionally similar INAs. The procedure was as follows: (1) randomly select an INA (denoted **R**) as an initial prototype representative; (2) form a provisional cluster containing all INAs whose distance from **R** is less than 0.5; (3) within this provisional cluster, identify the INA with the lowest average distance to all other INAs and designate it as the updated prototype representative (**R\***); (4) if **R\*** differs from **R**, update **R** = **R\*** and return to step 2. Otherwise, the cluster is considered converged, saved, and the algorithm proceeds to step 5; (5) repeat steps 1-4 while unassigned INAs remain as candidate prototype representatives; (6) if clusters share INAs, overlapping INAs are assigned to the cluster with the lower average pairwise distance.

#### Visualizations

Plots were constructed in Python using Matplotlib (v3.9.2). Spatial activity plots were derived from corresponding segmented LR-MoNNet images, with each spheroid’s segmented region filled according to the normalized time of its burst onset. Stitched raster plots were created by identifying segments in the burst raster that contained INAs of interest; however, instead of showing the full duration of bursts, scatter points were used to denote burst onsets. UMAP plots were generated by computing embeddings over pairwise distances between all INAs in the recordings using the umap (umap-learn v0.5.7) Python module. The resulting 2D embeddings were colored according to cluster identity or experimental condition. INA movies were created using the Matplotlib (v3.9.2) animation module, with each frame generated from the segmented MoNNet image in which each spheroid’s segmented region was assigned a value of 0 if not bursting, 1 if newly bursting, and 2 otherwise. Venn diagrams were created using the matplotlib-venn (v1.1.2) module. The Ca²⁺ imaging supplementary movies were generated using ImageJ (v1.54p). An average projection image across the time stack was computed. The average image intensity was scaled down by multiplying by 0.85 and then subtracted from the raw time-series data stack.

## DATA AVAILABILITY

The main data supporting the results in this study are available within the main Manuscript, Supplementary Figures and detailed Supplementary Videos. All the raw data will be made available on request.

## ACKNOWLEDGEMENTS

We thank all the members of the Tomer Lab for fruitful discussions and support. This work was supported by Research in Science and Engineering (RISE; Columbia University) award, Roy and Diana Vagelos Precision Medicine pilot award, and Columbia University Arts and Sciences startup grant to R.T.. A.A. was supported by fellowship from the Scholarship Office (SCO), United Arab Emirates. This work was partially supported by the Stavros Niarchos Foundation (SNF).

## AUTHOR CONTRIBUITONS

R.T. conceptualized the LR-MoNNet approach. C.G. and R.T. developed the IncStim hardware, and Y.C. implemented the software control. C.G. and R.T. developed the LR-MoNNet Ca²⁺ imaging and optical stimulation assays, and C.G. performed mouse LR-MoNNet Ca²⁺ imaging, ketamine treatment and optogenetic stimulation with feedback from A.U.. E.D.D., with feedback from C.G., established the GCaMP/ChrimsonR co-expression protocol in LR-MoNNets and generated all the mouse LR-MoNNet samples. A.U. and R.T. developed the data analysis framework, prepared illustrations for Figures, and with input from C.G. based on raw data preprocessing, performed the data analysis. A.A., K.W.L., and R.T. established human LR-MoNNet protocols, and A.A. conducted the associated experiments. B.L., S.A.K., and J.A.G. provided cell lines, animal resources and contributed expert inputs. R.T. wrote the manuscript with input from all authors and supervised the project.

