## Supplementary Info for "In vitro reconstitution of hippocampal cell assemblies"

### Supplementary material

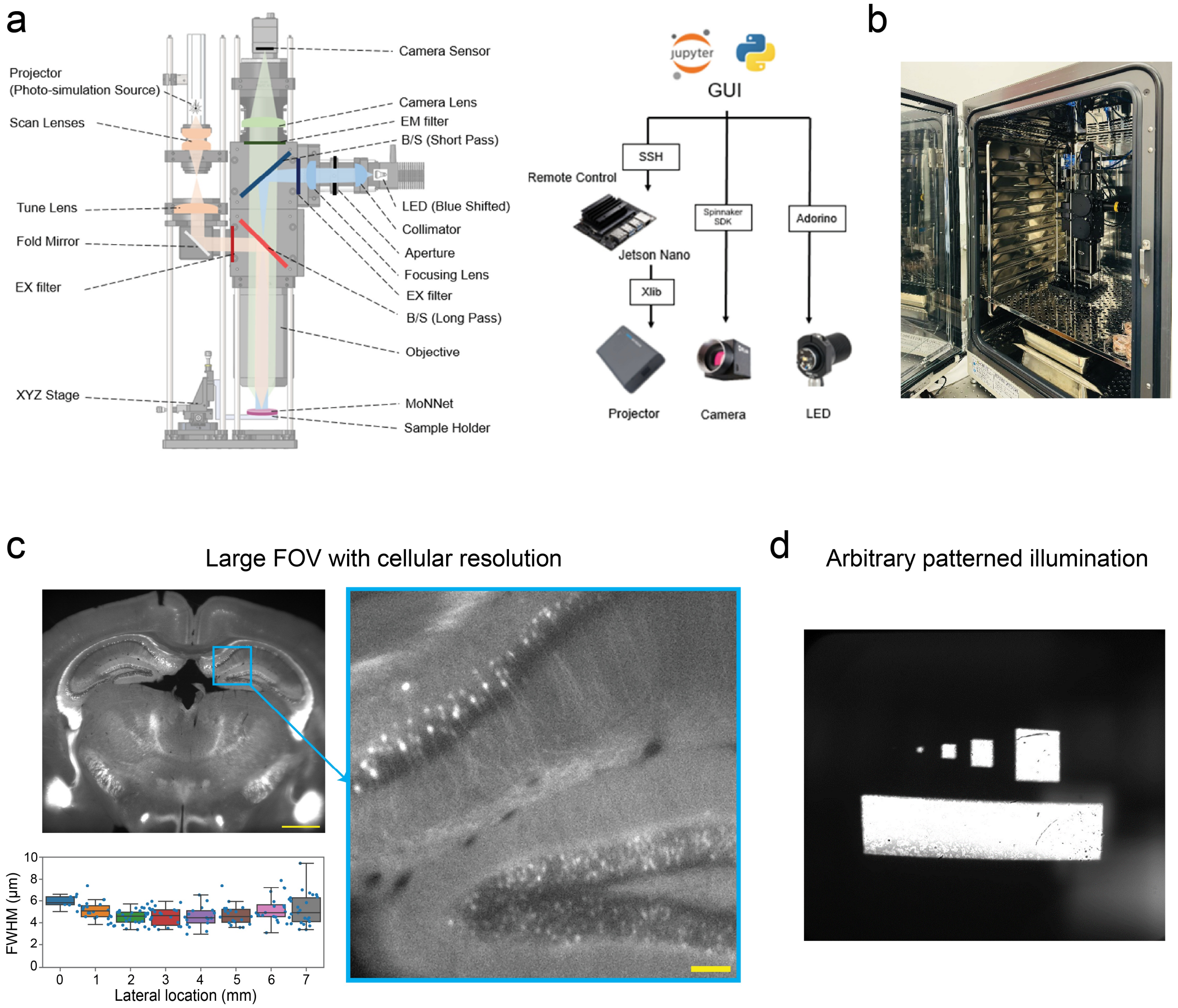


**Supplementary Figure 1 | IncStim Platform for All-Optical Interrogation of LR-MoNNets.** (**a**) *Left*: Schematic cross-section of the integrated imaging and stimulation optics. A projector delivers patterned optogenetic stimulation, while a blue-shifted LED provides wide-field excitation for $Ca^{2+}$ imaging. Excitation and emission paths are routed using excitation (EX) and emission (EM) filters and short-pass/long-pass dichroic beam splitters (B/S). Fluorescence is collected through the objective and imaged onto a camera sensor. Samples are mounted on a sample holder on an XYZ stage. *Right*: Control architecture for synchronized stimulation and imaging. A Python/Jupyter GUI communicates with the camera via the Spinnaker SDK, controls the projector through a Jetson Nano over SSH (projector interface via Xlib), and drives the LED via an Arduino. (**b**) Photograph of the fully assembled IncStim system mounted inside a standard tissue-culture incubator, illustrating the compact footprint and incubator-compatible form factor. (**c**) Large field-of-view imaging with cellular resolution. *Left*: Representative wide-field fluorescence image (FOV > 7 mm; cyan box indicates zoom region). *Right*: Zoomed view demonstrating cellular-scale features. *Bottom*: Lateral resolution across the field of view quantified as full width at half maximum (FWHM) versus lateral position. Scale bars: 1 mm (large view) and 100 µm (zoomed view). (**d**) Example of arbitrary patterned illumination projected onto the sample plane, demonstrating high-contrast patterns with sharp edges across multiple geometries.

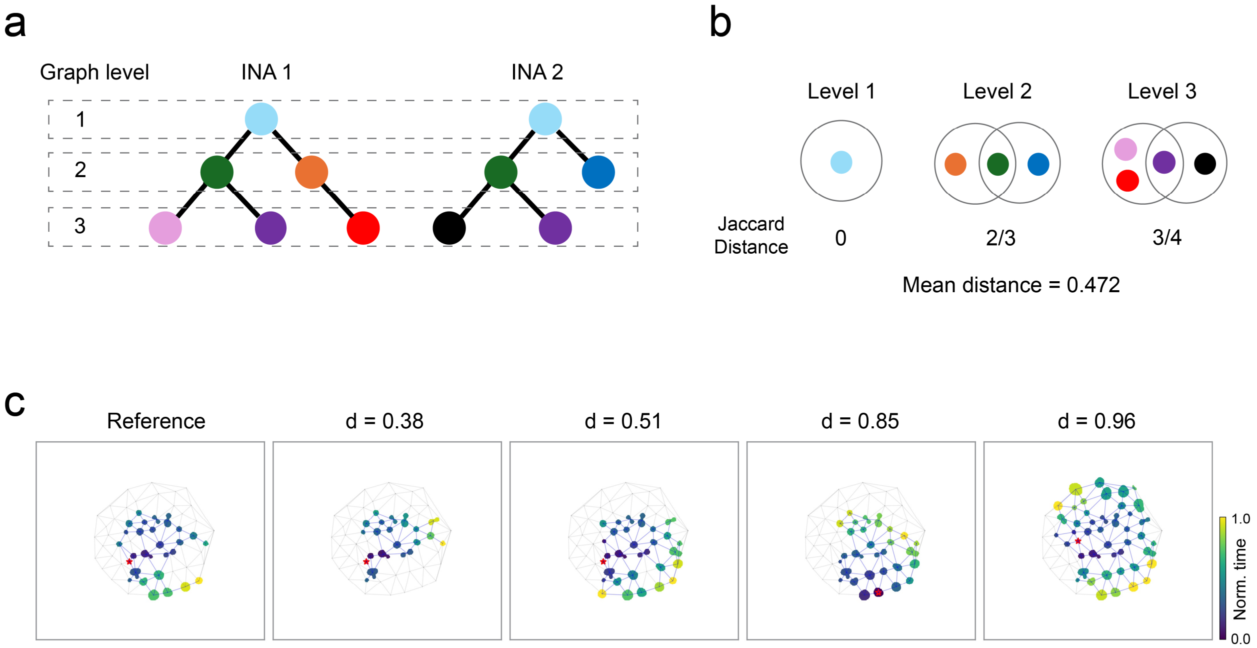


**Supplementary Figure 2 | Custom distance metric for INA clustering and analysis. (a)** Schematic representation of Intrinsic Network Assemblies (INAs) modeled as hierarchical graphs to define a custom distance metric capturing both spatial and temporal components of INA structure. **(b)** Illustration of the distance calculation between the two INAs shown in (a). To capture spatiotemporal similarity, the metric computes the Jaccard distance between the sets of active units at each corresponding graph level (temporal step), with the final distance defined as the mean of these level-specific distances. **(c)** Visualization of INA diversity relative to a "Reference" motif based on the calculated distance d. As the spatiotemporal trajectory of a candidate INA diverges from the reference, the distance value increases, validating the metric's utility for the clustering procedure used to identify cohesive sets of functionally similar INAs.

**Supplementary Table 1 | Bill of materials and component specifications for the IncStim all-optical platform.** This table lists all major optical, mechanical, electronic, and control components used to build the IncStim system. Together with **Supplementary Figure 1**, this table enables full reproducibility of the IncStim design using commercially available, off-the-shelf parts.

| Part | Item | Usage | Count | Price $ | Total |
|---|---|---|---|---|---|
| Stimulation arm | AnyBeam/Ultimems Projector AnyBeam Ultimems second generation (Sep/Oct 2023 launch) Ultimems LBS module | Laser source of Photo-stimulation | 1 | 200 | 200 |
| | Thorlabs: AC254-045-A | Plössl scan lens | 2 | 162 | 324 |
| | Thorlabs: AC508-075-A | Tube lens | 1 | 66 | 66 |
| | Thorlabs: CCM1-G01 | Fold mirror | 1 | 182 | 182 |
| | Thorlabs: DMSP567L | B/S long pass: stimulation | 1 | 653 | 635 |
| | Nvidia Jetson Nano | Operation of Laser source | 1 | 300 | 300 |
| | Thorlabs: MF630-69 | EX filter: ChR | | | |
| Illumination arm | Thorlabs:M455L4 | Blue shifted LED | 1 | 243 | 243 |
| | Thorlabs: LEDD1B | LED driver | 1 | 355 | 355 |
| | Thorlabs: ACL25416U-A | Collimator | 1 | 21 | 21 |
| | Thorlabs: CAB-LEDD1 | LED Connection Cable | 1 | 19 | 19 |
| | EO: 86-665 | EX filter: GCamp | 1 | 570 | 570 |
| | Thorlabs: CP20D | Aperture | 1 | 102 | 102 |
| | Thorlabs:AC254-100-A | Focusing lens | 1 | 89 | 89 |
| Imaging arm | Olympus: 1-SX965 1.6XPF | Objective | 1 | 2306 | 2306 |
| | EO: 66-253 | B/S short pass: imaging | 1 | 138 | 138 |
| | EO: 86-365 | EM filter: GCamp | 1 | 840 | 840 |
| | EO:49-284-INK | Camera lens | 1 | 190 | 190 |
| | Telednye Flir:BFS-U3-244S8M-C | Camera | 1 | 2619 | 2619 |
| Mechanics | VT 45N-25- XYZ-SK | XYZ Translation Stage | 1 | 1307 | 1307 |
| | Tube, cage, bars, bread board | Case system | N/A | 500 | 500 |
| Total | 11,006 | | | | |

**Supplementary Videos**:

**Supplementary Video 1 | Spontaneous activity sequences in Long-Range MoNNets.** Wide-field $Ca^{2+}$ imaging of two representative LR-MoNNet samples recorded (separately) using the IncStim platform inside an incubator. Spontaneous activity propagates sequentially across interconnected modular spheroids, revealing long-range structured dynamics spanning millimeter scales.

**Supplementary Video 2 | Chaining of Intrinsic Network Assemblies in Long-range MoNNets.** Representative $Ca^{2+}$ imaging dataset illustrating chaining of smaller intrinsic network assemblies (right columns) into a larger composite sequence (leftmost), analogous to *in vivo* hippocampal cell assembly dynamics. The top row shows $Ca^{2+}$ imaging data, and the bottom row show the corresponding detected INA motifs.

**Supplementary Video 3 | Resilience of intrinsic network assemblies (INAs) to optogenetic perturbation.** Spontaneous network activity recorded before, during, and after continuous optogenetic stimulation of approximately half of the LR-MoNNet sample (red rectangle). Despite prolonged perturbation, a subset of INAs re-emerges after the stimulation is off, demonstrating resilience to network-wide perturbations.

**Supplementary Video 4 | Pattern completion of an intrinsic network assembly following partial activation (example 1).** Four separate instances of targeted optogenetic activation of a single spheroid unit (red rectangle) trigger replay of an activation pattern closely matching a previously identified spontaneous INA, indicating attractor-like network dynamics in LR-MoNNets.

**Supplementary Video 5 | Pattern completion of an intrinsic network assembly across repeated trials (example 2).** Repeated optogenetic activation of the same spheroid unit over time reliably evokes the same assembly pattern. The stimulation window is indicated by the presence of a red rectangle on the target spheroid.

**Supplementary Video 6 | Acute ketamine-induced reconfiguration of intrinsic network assemblies.** $Ca^{2+}$ imaging of LR-MoNNets recorded at baseline, immediately after addition of 250 nM ketamine, one hour post-treatment, immediately after media washout, and one hour after washout. Ketamine exposure reconfigures the repertoire of active intrinsic network assemblies, with partial recovery following the washout.
